# A piezoelectric electroporator (Piezopen) for enhanced “naked” RNA vaccine delivery

**DOI:** 10.1101/2025.02.07.637103

**Authors:** Eleftheria Michalaki, Ananya Van Zanten, Jade Najjar, Gaurav Byagathvalli

## Abstract

Despite the success of COVID-19 mRNA vaccines, they still face challenges with high costs, complex manufacturing, off-target biodistribution, and systemic reactogenicity stemming from their inflammatory carriers: lipid nanoparticles (LNPs). While “naked” RNA delivery could in principle solve these issues, studies have suggested that it is infeasible due to rapid degradation by RNases and poor cellular entry, thereby necessitating formulations that enhance intracellular delivery and RNA stability. Now, we challenge this paradigm by showing that a simple and inexpensive (<$1), lighter-derived electroporator with microneedle electrodes (Piezopen) can augment gene expression and immunogenicity to naked mRNA leading to comparable responses to LNPs at low doses. We achieve robust responses in the absence of systemic inflammation and reactogenicity using skin-targeted delivery, administer diverse construct types (i.e., mRNA, self-amplifying RNA (saRNA), circular RNA (circRNA)), and demonstrate cross-species validation in live human skin to derisk subsequent clinical application. Our results introduce Piezopen as an inexpensive, well-tolerated, and efficacious alternative to LNPs for mRNA vaccine delivery, designed to facilitate routine vaccinations and pandemic response.

## 1. Introduction

mRNA vaccines have been revolutionary in terms of their rapid development and ability to prevent disease onset and severity, as exemplified during the COVID-19 pandemic.^1,2^ Despite their efficacy, they face challenges with expensive and complex manufacturing, excessive inflammation, poor reactogenicity, and limited durability. LNPs possess immunostimulatory properties, which provide potent adjuvant activity but also lead to significant side effects.^3^ In addition, these hurdles have impeded the uptake of mRNA vaccines in low- and middle-income countries, creating disparities in vaccine distribution. Addressing these challenges with LNPs and improving tolerability through an alternative delivery platform could facilitate further application of mRNA vaccines.

Localized delivery of “naked” RNA has been suggested as an approach to resolve issues with LNPs and other formulations while inducing immunity sufficient for protection.^2^ However, rapid degradation by RNases and poor intracellular uptake have previously resulted in poor responses even when delivering high payload doses. Successful application likely requires minimizing degradation via rapid and efficient intracellular delivery while inducing innate stimulation to adjuvant responses at doses that are not cost-prohibitive.

We previously described a platform that combines piezoelectric electroporation (EP) with microneedle (MN) electrodes (called ePatch or Piezopen) to efficiently deliver nucleic acid vaccines in the skin.^4^ This innovation precisely targets intracellular delivery in the immune-cell-rich epidermis while adjuvanting immune responses via localized cell death. The use of short, high-voltage electric pulses physically creates temporary pores in cell membranes enabling rapid uptake of nucleic acids. In previous studies, Piezopen improved gene expression and immune responses to naked DNA by >100-fold and >10-fold respectively, with superior tolerability compared to a hypodermic needle in humans.^5^

Building on our prior success^4^, we evaluated whether EP using Piezopen could deliver naked mRNA and induce immune responses comparable to LNPs at low doses. To do so, we investigated Piezopen’s ability to drive intracellular delivery of naked RNA (including, mRNA, self-amplifying RNA (saRNA), and circular RNA (circRNA)) and boost immune responses to a naked SARS-CoV-2 mRNA vaccine. We benchmarked to state-of-the-art LNPs clinically approved for RNA therapies, namely LNPs containing SM102, ALC0315, or MC3.^6^ In addition, we evaluated our hypothesis around cross-species translatability. Our study challenges the paradigm that suggests vaccination with naked mRNA is not feasible, uncovering a new modality for the delivery of mRNA vaccines more broadly.

## 2. Results

### Piezopen Significantly Augments Gene Expression and Induces Immune Responses to Naked mRNA with Comparability to LNPs *In Vivo*

To test our hypothesis that Piezopen (**Fig. 1A**) leads to successful intracellular delivery of naked mRNA *in vivo* superior to intradermal (ID) injection alone and comparable to LNPs, we conducted reporter expression kinetics studies using luciferase. We benchmarked against two state-of-the-art LNP formulations containing SM102 and ALC0315.^6^ We demonstrated that Piezopen delivery of mRNA-Luc leads to comparable expression magnitude and improved durability/kinetics compared to LNPs in a dose-dependent manner (**Fig. 1B**). Notably, gene expression after Piezopen administration persisted at a level 10-to 100-fold higher on Day 14 compared to LNPs, potentially improving antigen availability and vaccine durability. Together, Piezopen: (1) successfully delivered naked mRNA intracellularly and (2) mitigated mRNA degradation concerns.

**Figure 1.**
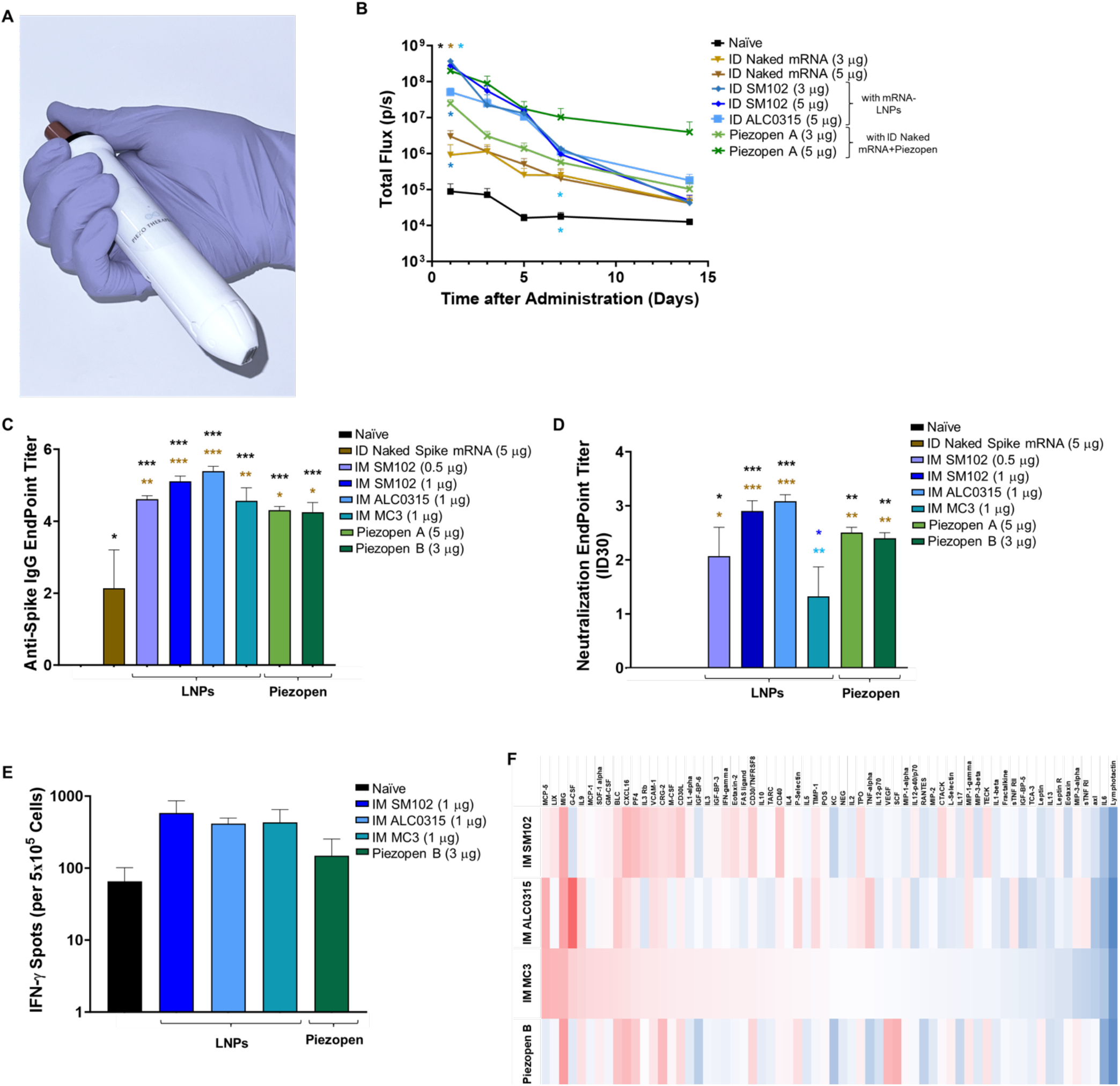
Piezopen augments gene expression and immune responses to naked RNA comparable to state-of-the-art LNPs. **(A)** Piezopen is an inexpensive (<$1), battery-free, handheld, and portable device that combines piezoelectric electroporation and microneedle electrodes. **(B)** Quantification of corresponding luciferase gene expression using IVIS. Total flux vs. time plot. **(C)-(E)** Quantification of binding, neutralizing antibodies, and IFN-γ spots for a SARS-CoV-2 mRNA vaccine using ELISA IgG, NAbs sVNT, and IFN-γ ELISpot kits. **(F)** Reactogenicity profile for LNPs and Piezopen assessed 24 hrs post-prime. Different doses (or routes of administration) are depicted by using different shades of the same color. *n*≥3/group, mean ± SEM, 2-way or 1-way ANOVA, and Tukey’s post hoc. \**p* < 0.05, \*\**p* < 0.01, \*\*\**p* < 0.001; color-coordinated asterisks above plots indicate a pairwise comparison.

After demonstrating we can augment intracellular delivery and gene expression to naked RNA with comparable magnitude to LNPs, we assessed whether Piezopen can induce immune responses to naked RNA vaccination. We benchmarked the delivery of a SARS-CoV-2 mRNA vaccine using Piezopen compared to three FDA-approved LNP formulations, quantifying humoral (anti-spike IgG and neutralizing antibody) and cellular (spike-specific IFN-γ production) responses along with reactogenicity. We selected doses between 0.5 µg and 1 µg for LNP groups to align with preclinical studies of approved COVID-19 vaccines while beginning with a higher dose of naked mRNA as initial proof-of-concept.^6^

We showed that Piezopen delivery of naked mRNA using Piezopen A (Piezopen design that led to the highest mRNA-Luc expression) induces robust immune responses significantly greater than the control conditions (both Naïve and ID injection alone) (**Fig. 1C-E**). In addition, manipulating EP field strength (Piezopen B) further augmented immune responses despite decreasing the mRNA dose by 40%, suggesting a viable path for optimizations to further reduce dosing while improving responses (Piezopen A: 5 µg vs Piezopen B: 3 µg). Delivery using both Piezopen A (5 µg) and Piezopen B (3 µg) led to comparable humoral and cellular immune responses to MC3 (1 µg) and slightly lower than SM102 (1 µg) and ALC0315 (1 µg). Importantly, neutralizing antibody titers induced by both Piezopen groups are similar to those reported in mRNA-1273 and BNT162b2 vaccines assessed using the same assay, suggesting clinical translatability.^7,8^

Quantification of a panel of relevant cytokines demonstrated an excellent systemic reactogenicity profile for all Piezopen groups and LNPs tested (**Fig. 1F**). Given the known inflammatory nature of LNPs, this is a rather surprising finding. Further cytokine evaluation in an earlier (4-6 h post-administration) and/or later (Day 22 following the booster) timepoint should be conducted in future studies.

Taken together, we challenge conventional wisdom by showing that Piezopen delivery of naked mRNA: (1) achieves comparable gene expression kinetics to benchmark LNPs, (2) induces robust humoral and cellular immune responses comparable to best-in-class LNP formulations, and (3) produces minimal reactogenicity concerns.

### Piezopen Significantly Augments Gene Expression to Diverse Array of Naked RNA Payloads with Comparability to LNPs Across Species *In Vivo* and in Live Human Skin *Ex Vivo*

By leveraging Piezopen across RNA and its subtypes, we can enable a diverse array of applications. Here, we explored Piezopen delivery of saRNA and circRNA, which have either shown enhanced antigen expression at lower doses (saRNA)^9–13^ or have been more stable and easier to store with lower innate immunogenicity, prolonged antigen-yielding capabilities, and durable immune responses (circRNA)^14,15^ compared to canonical linear mRNA. We validated saRNA delivery using reporters (i.e., luciferase) showing comparable expression magnitude and kinetics to state-of-the-art LNP (SM102) at an identical 1 µg dose (**Fig. 2A**). We showed that Piezopen delivery of naked circRNA leads to higher expression than ID injection alone, also at a low 1 µg dose (**Fig. 2B**). Our demonstration of naked mRNA/saRNA/circRNA delivery illustrates Piezopen can achieve comparable intracellular delivery and effectiveness to state-of-the-art LNP formulations in a “payload-agnostic” manner.

**Figure 2.**
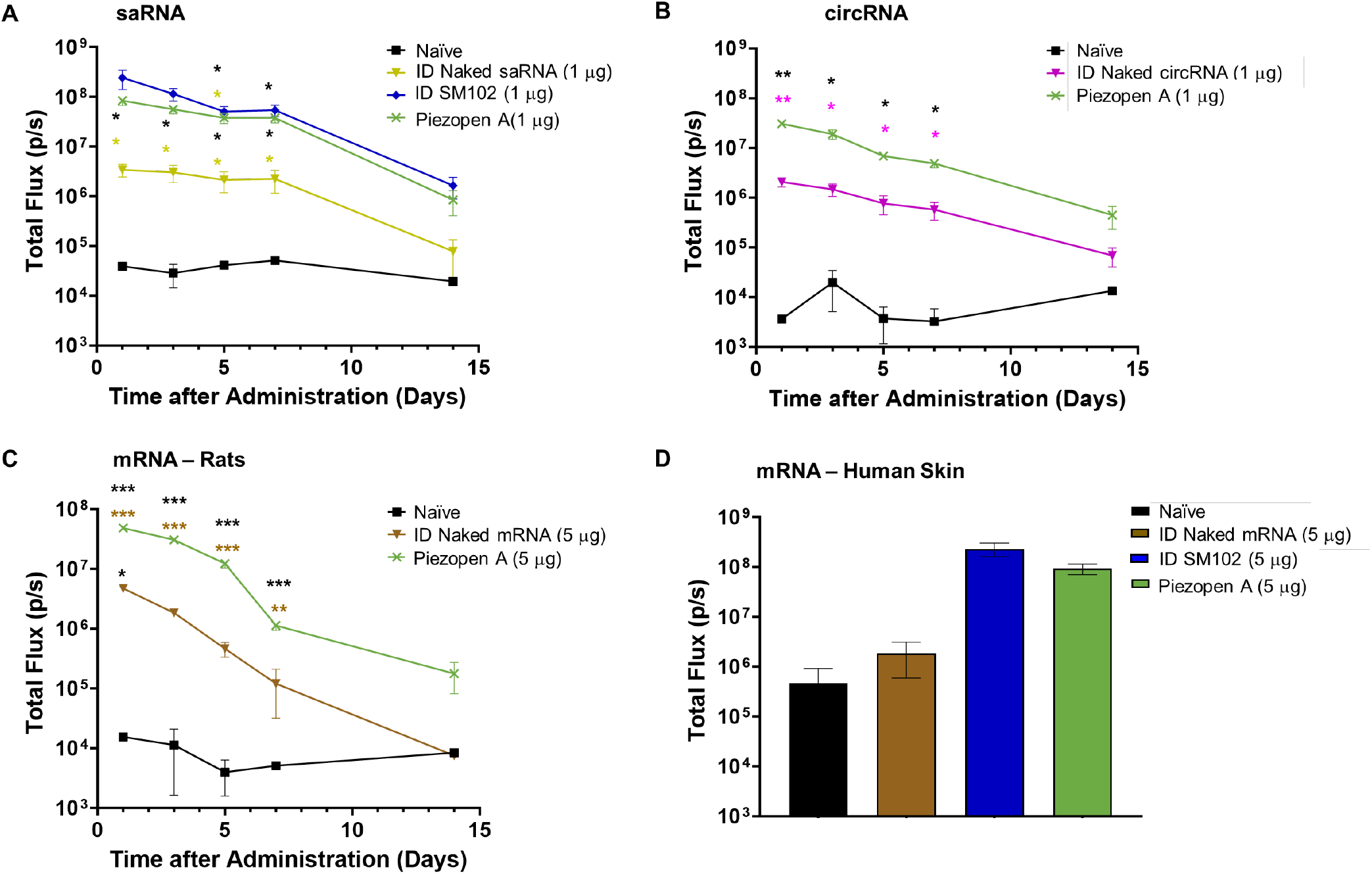
Piezopen drives intracellular delivery of naked RNA (namely, mRNA, saRNA, and circRNA) across species (namely, mice and rats) *in vivo* and in live human skin *ex vivo*. **(A)-(D)** Quantification of corresponding gene expression using IVIS via Total flux vs time plots. *n*≥3/group, mean ± SEM, 2-way or 1-way ANOVA, and Tukey’s post hoc. \**p* < 0.05, \*\**p* < 0.01, \*\*\*\**p* < 0.0001; color-coordinated asterisks above plots indicate a pairwise comparison.

To demonstrate our results are translatable across species, we evaluated mRNA delivery in rats and preserved human skin explants, the latter of which provides the closest available proxy to real-world human use. We show that Piezopen delivery significantly improves gene expression over injection alone in rats (**Fig. 2C**) and is comparable to SM102 LNP in human skin *ex vivo* (**Fig. 2D**), both at a low subtherapeutic dose (5 µg), indicating efficient intracellular delivery with minimal mRNA degradation. Since the Piezopen device used in the *ex vivo* human skin study is identical to what was used in mice, we confirm our results are translatable across species derisking clinical translation for human use. Lastly, comparable expression to LNPs at a low dose in human skin suggests Piezopen could potentially match or decrease clinical mRNA vaccine doses given the increased thickness and broader immune-cell repertoire of the human epidermis compared to mice.

## 3. Discussion

Our study reveals that naked RNA vaccination is feasible without LNPs or other formulation approaches currently considered necessary for cargo (RNA) protection and intracellular delivery^6,8^, presumably because the EP-based mechanisms deliver RNA into cells within seconds.^16^ Piezopen achieves comparable immunogenicity and improved expression kinetics compared to LNPs at low doses. This is a key finding as antigen persistence has been shown to significantly impact vaccine durability.^17,18^ Piezopen showed no reactogenicity concerns in this study despite using higher doses compared to LNPs, enabling multivalent or combination vaccines via increased tolerability/dosing thresholds. Piezopen delivery covers diverse payload sizes from ∼1,000 bp mRNA to ∼10 kbp saRNA/DNA, showcasing vast cargo capacities to deliver larger, complex, or multiple antigens. Notably, Piezopen induces robust immune responses to M1Ψ-modified linear mRNA which is the least stable and immunogenic platform^19^, indicating significant promise with circRNA and saRNA modalities. These findings uncover the benefits of Piezopen in improving mRNA vaccine durability via antigen persistence and adjuvantation, addressing a major limitation of mRNA vaccines.

The development of efficacious mRNA medicines is hindered by the limited understanding of the exact cell types that are necessary targets for effective protection. Studies have suggested that antigen-presenting cell (APC) transfection efficiency is associated with antigen-specific immune responses^20^, while non-APCs have been shown to play a significant role in inducing broad and robust immune responses relative to expression in APCs alone with other physical delivery systems.^21–24^ These studies suggest that cross-presentation by non-APCs to dendritic cells, rather than direct presentation by dendritic cells, is responsible for inducing the most robust humoral and cellular immune responses. LNP delivery targets immune cell types including T cells, natural killer (NK) cells, macrophages, and dendritic cells at the injection site (muscle) but also in the draining lymph nodes, spleen, and liver. In contrast, Piezopen concentrates pulses (and antigen expression) to the epidermis, targeting resident APCs, keratinocytes, Langerhans cells, etc. to maximize immunogenicity without off-target biodistribution.

Nonetheless, the observed differences in immune responses between Piezopen and LNPs could be attributed to the distinct cell populations and/or the total number of APCs transfected by each delivery mechanism.

While future research will further elucidate mechanism(s) by which Piezopen delivery achieves robust immune responses, we hypothesize a combination of efficient intracellular delivery, prolonged antigen expression, localized immune cell transfection, and adjuvantation via cell death^26^ drive innate and adaptive immunity. This study serves as a proof-of-concept toward successful naked RNA intracellular delivery and vaccination, and our findings further uncover potential benefits of Piezopen in improving RNA vaccine: 1) durability via antigen persistence and adjuvantation, and 2) tolerability via localized administration vs systemic uptake, addressing major limitations of LNP-delivered RNA vaccines.

Overall, our findings underscore Piezopen as a potent, inexpensive, and versatile delivery platform to facilitate increased acceptability and global translation of mRNA vaccines.

## 4. Acknowledgments

We thank M.R. Prausnitz, M.S. Bhamla, P. Rohilla, and C.L. Sundell for their helpful suggestions. We thank Kensei Komatsu and Jian-Dong Li for generously allowing us to use their IVIS system.

## 5. Materials and Methods

### Animal Studies

Female 6–8-week-old Balb/c mice or ∼8-week-old Wistar rats were used for the *in vivo* studies.

For immunogenicity studies, blood collection was performed via the lateral tail vein at various time points and processed for serum. During terminal procedures, blood was collected via a cardiac puncture or inferior vena cava and spleens were harvested.

All procedures were approved by the Georgia State Institutional Animal Care and Use Committee (IACUC) Review Board. Studies were carried out at the Division of Animal Resources (DAR), Georgia State University, Atlanta, GA.

### *Ex vivo* Human Skin Studies

Live human skin samples were purchased from Genoskin and maintained using the manufacturer’s protocol. Genoskin utilizes donated surgical human skin biopsies and stabilizes them in a proprietary matrix to preserve tissue viability and live skin responses for 7 days.

### Piezopen Administration

Piezopen consists of a piezoelectric electroporator coupled with microneedle electrodes that localize electric fields to the epidermis, as described previously.^4,5^ Briefly, using Piezopen involves performing a 20 µL intradermal (ID) injection via the Mantoux method (insulin syringe) followed by Piezopen application of 5 or 10 pulses. Bleb formation served as visual confirmation for a successful ID injection. For naked RNA injections, payloads were prepared by adding an RNase inhibitor to Ca^2+^-free PBS as described previously.^27^

### Reporter Expression Kinetics Studies

Animals (mice and rats) and human skin samples were injected with 5 µg FLuc-mRNA (SC2325, GenScript). For ID injections, a volume of 20 µL was used while for IM injections a volume of 50 µL. We benchmarked to FDA-approved LNP formulations, namely SM102, ALC0315, and MC3 (LNP-formulated mRNA and saRNA were purchased from GenScript).^6^ LNPs were administered ID for reporter studies. Mice were injected with 1 µg FLuc-saRNA (SC2346, GenScript) and 1 µg FLuc-circRNA (SC2339, GenScript).

### *In Vivo* Imaging System (IVIS)

IVIS was used 24 hrs after FLuc-RNA delivery. For animal studies, luciferin was injected intraperitoneally while for human skin samples, luciferin was injected via ID administration at the injection site. Samples were incubated for 10-15 min before imaging. IVIS was performed using an IVIS® Spectrum Imaging System. For animal reporter kinetics studies, animals were monitored for 14 days post-administration. For human skin reporter kinetics studies, human skin samples were imaged 1 day post-administration.

Vaccination Studies. Spike mRNA (SC2346, Genscript) was delivered to mice using ID injections with or without Piezopen electroporation and IM injection of LNPs encapsulated with Spike mRNA (SM102, ALC0315, and MC3) (Day 0). Injections were performed on the left flank (for ID) or left quadricep (for IM) of each animal at different doses (0.5 µg, 1 µg, 3 µg, 5 µg) depending on the group. A booster dose was administered on Day 21.

### Blood and Tissue Collection

A baseline blood collection was performed for serum processing a day prior to prime dose injections (pre-bleed, Day -1). A second blood collection was performed for serum processing on Day 14. On Day 35, a final blood collection and necropsy were performed, collecting blood for serum processing and harvesting the spleen for testing. Animal weight was monitored throughout the study.

### ELISA IgG

Spike protein-specific IgG levels in Day 35 serum were measured using the Mouse Anti-SARS-CoV-2 Antibody IgG kit (RAS-T023, ACRO Biosystems) according to the manufacturer’s protocol. Endpoint titers were calculated as the highest dilution emitting an optical density greater than 4x background.

### Neutralizing Antibodies

Levels of neutralization antibodies in Day 35 serum were measured using the SARS-CoV-2 Surrogate Virus Neutralization Test (sVNT) Kit (L00847-A, GenScript) according to the manufacturer’s protocol. Of note, not all replicates in SM102 (1 µg) and ALC0315 (1 µg) achieved endpoint titers due to limited assay availability to run additional dilutions. Endpoint titers were calculated as the highest dilution emitting an inhibition rate exceeding 30% based on the manufacturer’s protocol.

### ELISpot

IFN-γ ELISPOT assays were performed on Day 35 using a commercial kit Mouse IFN-γ Single-Color ELISPOT (96-well, precoated, strip, CTL) following the manufacturer’s protocol. In brief, 500,000 cells (from spleens) were stimulated overnight (17-18 hrs) with 2 μg/mL Spike peptide (5823-45-01, InvivoGen). SARS-CoV-2 Spike Glycoprotein-crude (RP30020, GenScript) was used as a positive control at a concentration of 2 μg/mL.

### Reactogenicity Array

A mouse cytokine array was performed 1-day post-administration with serum according to the manufacturer’s instructions (Mouse Cytokine Array G3 kit, AAM-CYT-G3-4, RayBiotech). The array was run and the analysis was performed by RayBiotech.

### Statistical Analysis

For animal reporter studies, a two-way ANOVA with Tukey’s multiple comparisons test was performed. For human skin reporter studies, ordinary one-way ANOVA with Tukey’s multiple comparisons test was performed. For vaccination studies, a two-way ANOVA with Tukey’s multiple comparisons test was performed. Each data point corresponds to either an independent subject (animal or human skin sample) or the average of technical replicates as stated in the figure caption. Error bars indicate the corresponding standard error of the mean. Data were analyzed using GraphPad Prism 7 (GraphPad Software).

Reported *p* values are multiplicity adjusted to account for multiple comparisons. For all cases, significance was defined as *p* < 0.05 (*) or *p* < 0.01 (**), or *p* < 0.001 (***).

## References

[1] May, M. After COVID-19 successes, researchers push to develop mRNA vaccines for other diseases. Nat Med 27, 930–932 (2021).

[2] Pardi, N., Hogan, M. J., Porter, F. W. & Weissman, D. mRNA vaccines — a new era in vaccinology. Nat Rev Drug Discov 17, 261–279 (2018).

[3] Ndeupen, S., Qin, Z., Jacobsen, S., Bouteau, A., Estanbouli, H. & Igyártó, B. Z. The mRNA-LNP platform’s lipid nanoparticle component used in preclinical vaccine studies is highly inflammatory. iScience 24, 103479 (2021).

[4] Xia, D., Jin, R., Byagathvalli, G., Yu, H., Ye, L., Lu, C. Y., Saad Bhamla, M., Yang, C. & Prausnitz, M. R. An ultra-low-cost electroporator with microneedle electrodes (ePatch) for SARS-CoV-2 vaccination. Proc Natl Acad Sci U S A 118, e2110817118 (2021).

[5] Lu, C. Y., Rohilla, P., Felner, E. I., Byagathvalli, G., Azizoglu, E., Bhamla, M. S. & Prausnitz, M. R. Tolerability of a piezoelectric microneedle electroporator in human subjects. Bioeng Transl Med 9, e10662 (2024).

[6] Wang, L., Davis, P. B., Kaelber, D. C., Volkow, N. D. & Xu, R. Comparison of mRNA-1273 and BNT162b2 Vaccines on Breakthrough SARS-CoV-2 Infections, Hospitalizations, and Death During the Delta-Predominant Period. JAMA 327, 678–680 (2022).

[7] Tangsathapornpong, A., Nanthapisal, S., Pontan, K., Bunjoungmanee, P., Neamkul, Y., Boonyarangkul, A., Wanpen, S., Fukpho, W., Jitpokasem, S., Tharabenjasin, P. & Jaru-Ampornpan, P. Immunogenicity and Safety of the Third Booster Dose with mRNA-1273 COVID-19 Vaccine after Receiving Two Doses of Inactivated or Viral Vector COVID-19 Vaccine. Vaccines (Basel) 11, 553 (2023).

[8] Liu, Y., Sánchez-Ovando, S., Carolan, L., Dowson, L., Khvorov, A., Hadiprodjo, J., Tseng, Y. Y., Delahunty, C., Khatami, A., Macnish, M., Dougherty, S., Hagenauer, M., Riley, K. E., Jadhav, A., Harvey, J., Kaiser, M., Mathew, S., Hodgson, D., Leung, V., Subbarao, K., Cheng, A. C., Macartney, K., Koirala, A., Marshall, H., Clark, J., Blyth, C. C., Wark, P., Kucharski, A. J., Sullivan, S. G. & Fox, A. Comparative B cell and antibody responses induced by adenoviral vectored and mRNA vaccines against COVID-19. medRxiv 2023.06.02.23290871 (2023).

[9] Bloom, K., van den Berg, F. & Arbuthnot, P. Self-amplifying RNA vaccines for infectious diseases. Gene Therapy 2020 28:3 28, 117–129 (2020).

[10] Ballesteros-Briones, M. C., Silva-Pilipich, N., Herrador-Cañete, G., Vanrell, L. & Smerdou, C. A new generation of vaccines based on alphavirus self-amplifying RNA. Curr Opin Virol 44, 145–153.

[11] Cu, Y., Broderick, K., Banerjee, K., Hickman, J., Otten, G., Barnett, S., Kichaev, G., Sardesai, N., Ulmer, J. & Geall, A. Enhanced Delivery and Potency of Self-Amplifying mRNA Vaccines by Electroporation in Situ. Vaccines (Basel) 1, 367–383 (2013).

[12] Vogel, A. B., Lambert, L., Kinnear, E., Busse, D., Erbar, S., Reuter, K. C., Wicke, L., Perkovic, M., Beissert, T., Haas, H., Reece, S. T., Sahin, U. & Tregoning, J. S. Self-Amplifying RNA Vaccines Give Equivalent Protection against Influenza to mRNA Vaccines but at Much Lower Doses. Molecular Therapy 26, 446–455 (2018).

[13] Saraf, A., Gurjar, R., Kaviraj, S., Kulkarni, A., Kumar, D., Kulkarni, R., Virkar, R., Krishnan, J., Yadav, A., Baranwal, E., Singh, A., Raghuwanshi, A., Agarwal, P., Savergave, L., Singh, S., Pophale, H., Shende, P., Shinde, R. B., Vikhe, V., Karmalkar, A., Deshmukh, B., Giri, K., Deshpande, S., Bulle, A., Siddiqui, M. S., Borthakur, S., Tummuru, V. R., Rao, A. V., Shukla, D., Jain, M. K., Bhardwaj, P., Supe, P. D., Das, M. K., Lahoti, M. & Barge, V. An Omicron-specific, self-amplifying mRNA booster vaccine for COVID-19: a phase 2/3 randomized trial. Nature Medicine 2024 30:5 30, 1363–1372 (2024).

[14] Chen, R., Wang, S. K., Belk, J. A., Amaya, L., Li, Z., Cardenas, A., Abe, B. T., Chen, C. K., Wender, P. A. & Chang, H. Y. Engineering circular RNA for enhanced protein production. Nature Biotechnology 2022 41:2 41, 262–272 (2022).

[15] Liu, X., Zhang, Y., Zhou, S., Dain, L., Mei, L. & Zhu, G. Circular RNA: An emerging frontier in RNA therapeutic targets, RNA therapeutics, and mRNA vaccines. J Control Release 348v 84 (2022).

[16] Knudsen, M. L., Ljungberg, K., Liljeström, P. & Johansson, D. X. Intradermal electroporation of RNA. Methods Mol Biol 1121, 147–154 (2014).

[17] Joyce, J. C., Sella, H. E., Jost, H., Mistilis, M. J., Esser, E. S., Pradhan, P., Toy, R., Collins, M. L., Rota, P. A., Roy, K., Skountzou, I., Compans, R. W., Oberste, M. S., Weldon, W. C., Norman, J. J. & Prausnitz, M. R. Extended delivery of vaccines to the skin improves immune responses. J Control Release 304, 135 (2019).

[18] Palin, A. C., Alter, G., Crotty, S., Ellebedy, A. H., Lane, M. C., Lee, F. E. H., Locci, M., Malaspina, A., Mallia, C., McElrath, M. J., Pulendran, B., Singh, A. & D’Souza, M. P. The Persistence of Memory: Defining, Engineering, and Measuring Vaccine Durability. Nat Immunol 23, 1665 (2022).

[19] Mokuda, S., Watanabe, H., Kohno, H., Ishitoku, M., Araki, K., Hirata, S. & Sugiyama, E. N1-methylpseudouridine-incorporated mRNA enhances exogenous protein expression and suppresses immunogenicity in primary human fibroblast-like synoviocytes. bioRxiv 2022.03.22.485393 (2022).

[20] Chen, J., Xu, Y., Zhou, M., Xu, S., Varley, A. J., Golubovic, A., Ze Lu, R. X., Wang, K. C., Yeganeh, M., Vosoughi, D. & Li, B. Combinatorial design of ionizable lipid nanoparticles for muscle-selective mRNA delivery with minimized off-target effects. Proc Natl Acad Sci U S A 120, e2309472120 (2023).

[21] Lauterbach, H., Gruber, A., Ried, C., Cheminay, C. & Brocker, T. Insufficient APC Capacities of Dendritic Cells in Gene Gun-Mediated DNA Vaccination. The Journal of Immunology 176, 4600–4607 (2006).

[22] Cho, J. H., Youn, J. W. & Sung, Y. C. Cross-priming as a predominant mechanism for inducing CD8(+) T cell responses in gene gun DNA immunization. The Journal of Immunology 167, 5549–5557 (2001).

[23] Vij, R., Lin, Z., Schneider, K., Seshasayee, D. & Koerber, J. T. Analysis of the effect of promoter type and skin pretreatment on antigen expression and antibody response after gene gun-based immunization. PLoS One 13, e0197962 (2018).

[24] Kim, B. S., Miyagawa, F., Cho, Y. H., Bennett, C. L., Clausen, B. E. & Katz, S. I. Keratinocytes Function as Accessory Cells for Presentation of Endogenous Antigen Expressed in the Epidermis. Journal of Investigative Dermatology 129, 2805–2817 (2009).

[25] Kim, E. H., Teerdhala, S. V., Padilla, M. S., Joseph, R. A., Li, J. J., Haley, R. M. & Mitchell, M. J. Lipid nanoparticle-mediated RNA delivery for immune cell modulation. Eur J Immunol 2451008 (2024).

[26] Batista Napotnik, T., Polajžer, T. & Miklavčič, D. Cell death due to electroporation – A review. Bioelectrochemistry 141, 107871 (2021).

[27] Huysmans, H., De Temmerman, J., Zhong, Z., McCafferty, S., Combes, F., Haesebrouck, F. & Sanders, N. N. Improving the Repeatability and Efficacy of Intradermal Electroporated Self-Replicating mRNA. Mol Ther Nucleic Acids 17, 388–395 (2019).

